# Caspase-mediated regulation and cellular heterogeneity of the cGAS/STING pathway in Kaposi’s sarcoma-associated herpesvirus infection

**DOI:** 10.1101/2021.05.03.442439

**Authors:** Tate Tabtieng, Rachel C. Lent, Machika Kaku, Alvaro Monago Sanchez, Marta Maria Gaglia

**Author notes:** Correspondence to: Marta Maria Gaglia. These authors contributed equally (Tate Tabtieng, Rachel C. Lent).

## Abstract

As a result of the ongoing virus-host arms race, viruses have evolved numerous immune subversion strategies, many of which are aimed at suppressing the production of type I interferons (IFNs). Apoptotic caspases have recently emerged as important regulators of type I IFN signaling in both non-infectious contexts and during viral infection. Despite being widely considered anti-viral factors since they can trigger cell death, several apoptotic caspases promote viral replication by suppressing innate immune response. Indeed, we previously discovered that the AIDS-associated oncogenic gammaherpesvirus Kaposi’s sarcoma-associated herpesvirus (KSHV) exploits caspase activity to suppress the antiviral type I IFN response and promote viral replication. However, the mechanism of this novel viral immune evasion strategy is poorly understood, particularly how caspases antagonize IFN signaling during KSHV infection. Here we show that caspase activity inhibits the DNA sensor cGAS during KSHV lytic replication to block type I IFN induction. Furthermore, we use single-cell RNA-sequencing to reveal that the potent antiviral state conferred by caspase inhibition is mediated by an exceptionally small percentage of IFN-β-producing cells, thus uncovering further complexity of IFN regulation during viral infection. Collectively, these results provide insight into multiple levels of cellular type I IFN regulation that viruses co-opt for immune evasion. Unraveling these mechanisms can inform targeted therapeutic strategies for viral infections and reveal cellular mechanisms of regulating interferon signaling in the context of cancer and chronic inflammatory diseases.

**Importance:** Type I interferons are key factors that dictate the outcome of infectious and inflammatory diseases. Thus, intricate cellular regulatory mechanisms are in place to control IFN responses. While viruses encode their own immune-regulatory proteins, they can also usurp existing cellular interferon regulatory functions. We found that caspase activity during lytic infection with the AIDS-associated oncogenic gammaherpesvirus Kaposi’s sarcoma-associated herpesvirus inhibits the DNA sensor cGAS to block the antiviral type I IFN response. Moreover, single-cell RNA-sequencing analyses unexpectedly revealed that an exceptionally small subset of infected cells (<5%) produce IFN, yet this is sufficient to confer a potent antiviral state. These findings reveal new aspects of type I IFN regulation and highlight caspases as a druggable target to modulate cGAS activity.

## Introduction

Type I interferon (IFN) cytokines are potently anti-viral and play an essential role in controlling infections and starting systemic immune responses. Type I IFNs are generally turned on during viral infection after sensing of viral nucleic acids by cellular pattern recognition receptors such as the cytosolic DNA sensor cGAS and RNA sensor RIG-I(1). These sensors signal through adaptor proteins (STING and MAVS, respectively) and the kinases TBK1 and IKKε to activate the transcription factor IRF3 and trigger induction of type I IFN cytokines(1). Type I IFNs in turn stimulate expression of interferon stimulated genes (ISGs) in an autocrine and paracrine manner to limit viral replication and infection, as well as alert the immune system of the infection(2). As ISGs create a strong antiviral response, all viruses have evolved one or more strategies to block IFN induction(3). The AIDS-associated oncogenic gammaherpesvirus Kaposi’s sarcoma-associated herpesvirus (KSHV), which is the etiological agent for one of the leading causes of cancer death in sub-Saharan Africa, Kaposi’s sarcoma, is a prime example of this. KSHV induces minimal type I IFN responses when reactivating from latency(4, 5), demonstrating it can strongly block this response.

While many viral encoded factors directly target the type I IFN response, viruses like KSHV have also evolved strategies to usurp cellular pathways to counteract immune responses. We recently established that KSHV exploits the activity of the apoptotic caspases to suppress the type I IFN response and promote viral replication(4). Indeed, while KSHV lytic reactivation normally does not induce a strong type I IFN response, lytically infected cells express and secrete high levels of IFN-β if caspases are inhibited(4). Although apoptotic caspases are canonically thought to be antiviral by facilitating cell death of infected cells, multiple recent studies have demonstrated caspase-dependent inhibition of innate immune signaling during apoptosis and viral infection(4, 6–8). However, it remained unclear from our previous studies how caspases enact this IFN regulatory function during KSHV infection. In particular, while caspase-3 and -7 can directly cleave cGAS, MAVS, and IRF3 to block type I IFN signaling(8), our previous findings suggest that in the case of KSHV infections, caspase-8 is the major caspase suppressing the type I IFN response(4). Moreover, caspase-8 appears to act independently of caspase-3/7 activation and of cell death to regulate IFNs in KSHV-infected cells(4). As different caspases have different proteolytic targets, these results suggest the mechanism we found in KSHV infected cells may also be different than what has previously been reported.

To decipher how caspases disrupt IFN signaling during KSHV infection, we used RNAi-mediated depletion and pharmacological inhibition to identify which IFN signaling components are targeted by caspase activity. This revealed that caspase activity specifically inhibits cGAS to impede IFN signaling during KSHV lytic infection. We also performed single-cell RNA-sequencing (scRNAseq) to examine the heterogeneity of immune responses and viral gene expression during KSHV lytic infection. This uncovered considerable heterogeneity of the KSHV lytic cycle. Moreover, it surprisingly revealed that a small percentage of infected cells produce IFN and suggested that differences in cellular gene expression may account for this heterogeneity.

## Results

### Caspase activity blocks cGAS/STING signaling during KSHV lytic infection

We previously showed that the type I IFN response that occurs upon caspase inhibition in KSHV-infected cells is dependent on the kinase TBK1(4). TBK1 can be activated by several pathogen sensors, including sensors of viral nucleic acids(1), and both the DNA and the RNA sensing pathways can be activated during KSHV infection(5, 9–15). To discern which pathogen sensing pathway is inhibited by caspase activity during KSHV infection, we knocked down key proteins for DNA and RNA sensing: the DNA sensors cGAS and IFI16 and their signaling adaptor STING, and the RNA sensor RIG-I and its signaling adaptor MAVS (Fig. 1A-C, Supp Fig. 1). For these experiments, we used iSLK.219 cells, which are latently infected with recombinant rKSHV.219. rKSHV.219 expresses GFP constitutively and RFP under the control of a KSHV lytic promoter(16). These cells also express an exogenous copy of the KSHV master lytic regulator RTA under a doxycycline inducible promoter, so that reactivation of the viral lytic replication cycle can be induced by doxycycline treatment(16). IFN-β induction upon caspase inhibition was severely reduced by STING and cGAS depletion, suggesting that caspase activity targets STING-dependent type I IFN signaling during KSHV replication (Fig. 1A). Moreover, upon STING and cGAS knock-down, caspase inhibitors no longer caused a reduction in KSHV replication, which we previously showed is a result of type I IFN induction(4) (Fig.1B). In contrast, IFI16, RIG-I, and MAVS knockdowns had minimal effects on IFN-β induction and viral gene expression in the presence of caspase inhibitors (Fig. 1A, B). These results suggest that cGAS/STING-mediated DNA sensing is disabled by caspase activity. To corroborate this conclusion, we also tested pharmacological inhibition of cGAS enzymatic activity with RU.521. When bound to DNA, cGAS synthesizes the cyclic dinucleotide 2’3’-cGAMP, which in turn binds and activates STING(17). Consistent with our knock-down results, cGAS inhibition reduced induction of IFN-β and of the interferon-stimulated gene ISG15 upon caspase inhibition (Fig. 1D). Collectively, these findings suggest that caspase activity prevents activation or signal transduction of the cGAS/STING DNA sensing pathway during KSHV lytic replication.

**Figure 1.**
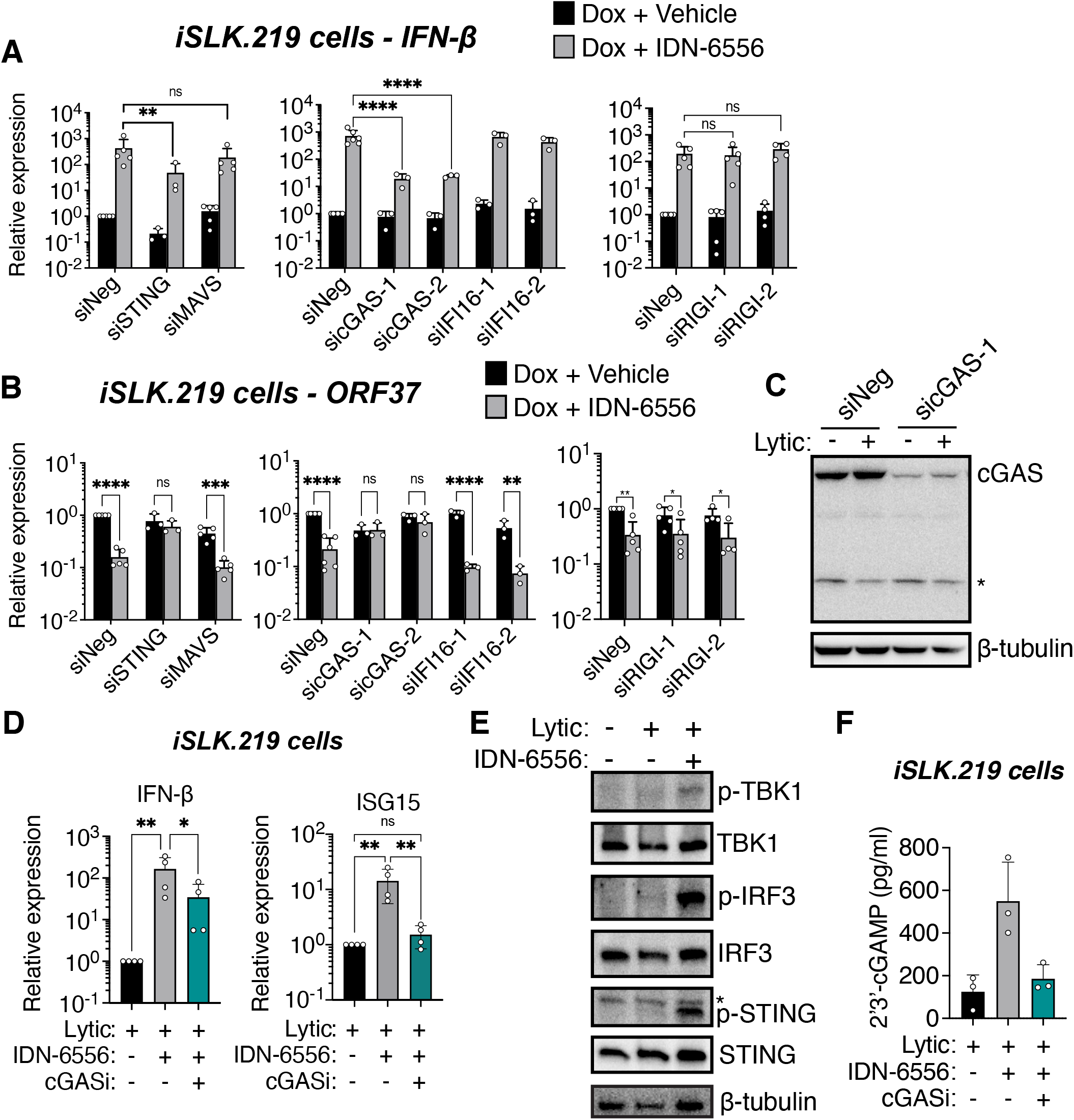
Caspase activity prevents cGAS activation during KSHV lytic replication to block IFN-β induction. (A-C) iSLK.219 cells were transfected with a negative control siRNA or siRNAs targeting the indicated proteins. For cGAS and RIG-I, the transfection was carried out twice, two days prior to and on the day of lytic cycle induction; for STING, MAVS and IFI16, one transfection was carried out two days prior to induction. The cells were then lytically reactivated with doxycycline (1 μ g/ml) and treated with either DMSO (vehicle) and IDN-6556 (10 μM) as indicated. (A-B) Total RNA was extracted at day 5 post reactivation and the levels of IFN-β (A) and ORF37 (B) mRNA were measured by RT-qPCR and normalized to levels of 18S rRNA (n ≥ 3). (C) Cell lysates were harvested at day 4 post reactivation and subjected to western blot for cGAS and β-tubulin as a loading control. Asterisk indicates non-specific band. Blots are representative of 3 replicates. (D-F) iSLK.219 cells were lytically reactivated with doxycycline (1 μg/ml) and treated with either DMSO (vehicle) or IDN-6556 (10 μM) and the cGAS inhibitor RU.521 (cGASi, 24.1 μM), where indicated. (D) Total RNA was extracted from iSLK.219 cells 3 days after reactivation. Levels of IFN-β and ISG15 mRNA were measured by RT-qPCR and normalized to 18S rRNA (n = 3). (E) Cell lysates were harvested at day 4 post reactivation and subjected to western blotting for p-TBK-1, TBK-1, p-IRF-3, IRF-3, p-STING, STING, and β-tubulin (as a loading control) as indicated. Asterisk indicates non-specific band.Blots are representative of 3 replicates. (F) Levels of 2’3’-cGAMP in lysate collected from iSLK.219 cells at day 4 post reactivation were measured by ELISA (n = 4). ns, *, **, ***, ****: p value > 0.05, < 0.05, < 0.01, < 0.001, < 0.0001, Tukey’s multiple comparisons test after two-way ANOVA.

### Caspases inhibit cGAS activity without reducing cGAS protein levels

To determine how caspases interfere with cGAS/STING signaling, we tested whether there was differential activation of various effectors in the cGAS/STING pathway in the presence *vs*. absence of caspase activity. We observed increased phosphorylation of the downstream proteins STING, TBK1, and IRF3, which indicates that they are activated upon caspase inhibition (Fig.1E). These results suggest that activation of the entire pathway is blocked by caspases. Indeed, we also found that 2’3’-cGAMP levels were increased upon caspase inhibition and were reduced by treatment with a cGAS inhibitor (Fig.1F). Taken together with our results from the knockdown experiments, these findings identify the target of caspases as a component or regulator of the cGAS/STING pathway that acts to promote activation or activity of cGAS.

We considered the possibility that the target was cGAS itself. One report suggests that caspase-3 and -7 can cleave cGAS during apoptosis and other viral infections(8). However, we have found that the activity of these caspases is dispensable for IFN regulation during KSHV lytic infection, and that caspase-8 is likely the key regulator(4). Also, we do not detect an overall reduction in cGAS levels nor any cGAS cleavage fragments during lytic reactivation (Fig. 1C). Ectopic co-expression of caspase-8 and cGAS also did not reveal caspase cleavage fragments, even though caspase-8 overexpression also leads to activation of caspase-3 and -7 (data not shown). It is thus likely that during KSHV lytic infection, caspases do not cleave cGAS directly, but instead target a regulatory factor that modulates its activity.

### Caspase activity also regulates cGAS in KSHV-infected primary effusion lymphoma B cells

To determine if caspases also suppress the cGAS/STING pathway in a cell type that is physiologically relevant for KSHV infection and tumorigenesis, we examined the activity of caspases in BC3 cells, a KSHV-infected B cell line derived from a primary effusion lymphoma (PEL) patient (18). Similarly to iSLK.219 and BCBL1 cells(4), caspase-8 was also activated upon lytic reactivation of BC3 cells (Fig. 2A). Moreover, caspase inhibition increased expression of both IFN-β and ISGs (ISG15 and IFIT1, Fig. 2B), as well as IRF3 phosphorylation (Fig. 2C). Consistent with what we observed in iSLK.219 cells, caspase inhibition potentiated 2’3’-cGAMP production in lytically reactivated BC3 cells (Fig. 2D). Moreover, inhibition of cGAS with RU.521 reduced the induction of IFN-β (Fig. 2E). These results strengthen our conclusion that caspases have a major role in inhibiting cGAS activity or activation to and blocking IFN induction during KSHV lytic infection in multiple cell types, including cell types that are relevant to KSHV infection in humans and KSHV-associated diseases.

**Figure 2.**
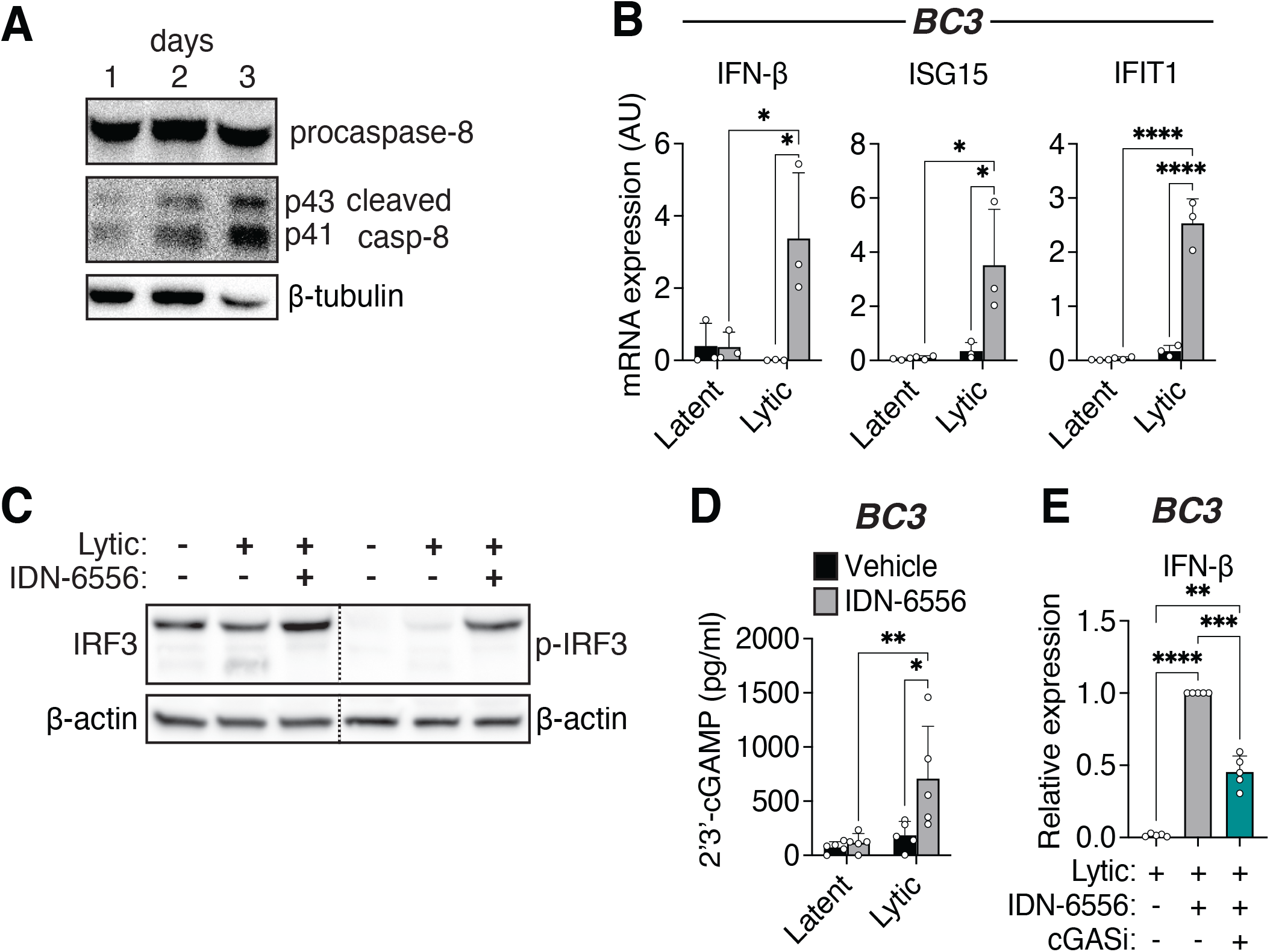
Caspases also inhibit cGAS activity in KSHV infected B cells. BC3 cells were lytically reactivated with TPA (20 ng/ml) for 1-3 days as indicated (A) or for 2 days (B-E) and treated with DMSO (vehicle), the pan-caspase inhibitor IDN-6556 (10 μM), and/or the cGAS inhibitor RU.521 (cGASi, 24.1 μM) as indicated. (A) Cell lysates were collected from latent and lytically reactivated BC3 cells at the indicated time points and subjected to western blot analysis for activated (cleaved) caspase-8, and β-tubulin as a loading control. Blots are representative of 3 replicates. (B) Total RNA was extracted from BC3 cells. Levels of the indicated mRNAs were measured by RT-qPCR and normalized to 18S rRNA (n = 3). (C) Cell lysates were collected from lytically reactivated BC3 cells and subjected to western blot analysis for activated p-IRF3, total IRF3, and β-actin as a loading control. The lower band for total IRF3 is likely a caspase-3 product as described by Aresté et al.52 Blots are representative of 3 replicates. (D) Levels of 2’3’-cGAMP in lysate collected from BC3 were measured by ELISA (n = 5). (D) Total RNA was extracted from BC3 cells. Levels of IFN-β mRNA were measured by RT-qPCR and normalized to 18S rRNA (n = 5). *, **,***,****: p value < 0.05, < 0.01, < 0.001, < 0.0001, Tukey’s multiple comparisons test after two-way ANOVA.

### Single-cell RNA sequencing (scRNA-Seq) highlights heterogeneity of KSHV reactivation and confirms the pro-viral role of caspase activity

The results so far provide a bulk analysis of the IFN pathway during KSHV infection. However, not all iSLK.219 cells in the population re-enter the lytic cycle upon doxycycline addition(16), suggesting there could be significant cell-to-cell variability in responses to infection. To understand both lytic reactivation and IFN signaling on a single cell basis, we carried out scRNA-Seq on uninfected cells, latently infected cells, lytically infected cells, and lytically infected cells treated with IDN-6556 (Fig. 3A, Table S1). Uninfected cells are SLK cells containing the inducible KSHV RTA transgene, but no rKSHV.219 (iSLK.RTA)(16). We also included lytically infected cells treated with IDN-6556 and anti-IFN antibodies to block autocrine and paracrine signals (Fig. 3A, Supp. Fig. 2A, Table S1). Analysis using the Seurat package(19, 20) unbiasedly identified 13 clusters with different gene expression across the 4 samples (Fig. 3B). Analysis of the mapped reads showed that, as expected, there was a higher percentage of KSHV reads in cells that were lytically reactivated (Fig. 3C, D). Certain clusters were clearly enriched in KSHV transcripts (Fig.3C, E; Supp. Fig. 2B) and these corresponded to clusters that only appeared in the lytic samples (Fig. 3B, C). Analysis of the cell cycle status indicated that most of the cells were in G1, but some of the clusters were enriched in cells in S or G2/M phase, and this tended to correlate with lower viral gene expression, presumably indicative of lower reactivation (Supp. Fig. 2C). We further analyzed the type of genes that were seen in the different clusters in the lytic+vehicle sample. We saw that several clusters had cells with on average 5-30% of reads mapping to KSHV genes, while more than 75% of the reads mapped to KSHV genes in cells from cluster 12 (Fig. 3E-F, Supp. Fig 2B). Because our scRNA-Seq is directed at the 3’ end of RNAs and many mRNAs in KSHV have the same 3’ end(21, 22), we were not able to conclusively distinguish expression of all the viral genes. Therefore, in the heatmaps we list together all the ORFs that each 3’ end corresponds to. Nonetheless, analysis of viral genes and of the rKSHV.219-encoded GFP, RFP and puromycin-N-acetyl transferase (PAC) reporters agrees with the general picture obtained from percentage reads. Most clusters show some lytic gene expression but only a few demonstrate substantial expression of lytic genes (Fig. 3E, Supp. Fig. 2B). Interestingly, we note that some of the clusters that expressed lytic genes did not express high levels of RFP (cluster 0 for example, Fig. 3E). Because KSHV lytic genes are expressed in several waves, termed early lytic, delayed early and late(23–25), we roughly categorized the clusters based on the pattern of lytic gene expression, as well as the cell cycle stage and the expression of IFNs. We classified clusters as “uninfected”, “latent”, “latent (dividing)”, “lytic (early)”, “lytic (intermediate)”, “lytic (late)”, “lytic (early) and IFN+” (Fig. 3E, 3G-H, Fig. 4A-C). Most of the viral transcripts detected increased in levels monotonically in cells that had progressed further through the lytic cycle, except for ORF57, K4.1, and vIL-6, which appeared to decrease in levels in the “late” lytic cell cluster (cluster 12, Figure 3E). Figure 3G and Supp. Fig. 3B show how the different clusters expanded and contracted in each sample. Interestingly, addition of the caspase inhibitors and the consequent type I IFN induction did not reduce the overall percentage of cells expressing lytic genes (Fig. 3G, Table S2), but it reduced the number of cells that we classified as “lytic (intermediate)” and “lytic (late)” (particularly clusters 7, 8, and 12) (Fig. 3G). Moreover, caspase inhibitor treatment reduced the levels of most viral genes in many clusters, most notably the “lytic” clusters 5, 8, and 12 (Fig. 3E, H, Supp. Fig. 2B, Supp. Fig. 3). The effect of caspase inhibitors on the size of these clusters and the reduction in lytic gene expression was largely rescued by blocking type I IFN signaling with anti-IFN antibodies (Fig. 3G, H, Supp. Fig. 2B, Supp. Fig. 4). These results confirm that caspase activity is required for reactivation and/or progression through the lytic cycle and that inhibition of the lytic cycle upon caspase inhibition is due to the paracrine and autocrine effects of type I IFN secretion. These data thus recapitulate our bulk level findings on the role of caspases during KSHV replication (4). The scRNA-Seq results also confirm the wide variability in stage of reactivation of iSLK.219 cells and reveal that the RFP signal in iSLK.219 cells may underestimate the cells that are expressing lytic genes.

**Figure 3.**
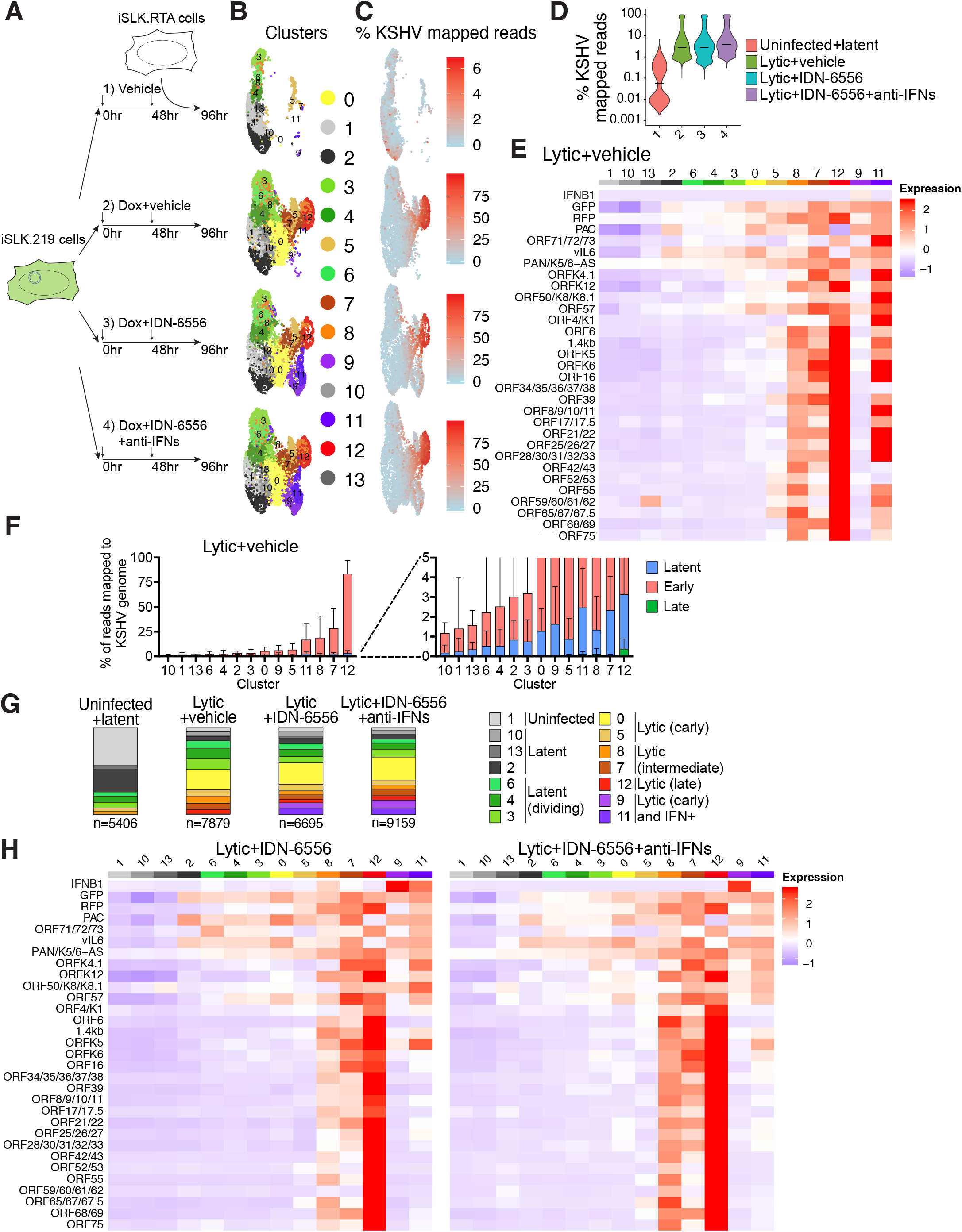
Single-cell RNA-Seq analysis reveals subsets of lytically reactivating cells. (A) Diagram of the single-cell RNA-Seq (scRNA-Seq) experiment using 4 samples. Sample 1 was a mixture of uninfected iSLK.RTA and latently infected iSLK.219 cells treated with DMSO (vehicle). Samples 2-4 were iSLK.219 treated with doxycycline (Dox) to reactivate lytic replication (“Lytic”) and treated with DMSO (vehicle), the caspase inhibitor IDN-6556, or IDN-6556 and a cocktail of anti-IFN antibodies to block paracrine IFN signaling. All samples were collected 4 days after the start of treatments. (B-H) Analysis of data from the scRNA-Seq experiment presented in (A). (B) Uniform manifold analysis and projection (UMAP) diagram of the clusters unbiasedly defined by Seurat in the 4 samples. (C) UMAP diagram showing what percentage of reads from each cell comes from the KSHV genome. Legends on the right define what percentage the colors represent. (D) Violin plot of the distribution of cells in each sample based on percentage of KSHV reads. Sample 1 has a bimodal distribution because it is a mix of uninfected and latent cells. (E) Heatmap of average expression of KSHV genes in each cluster in the lytic + vehicle sample. The legends on the right define what expression level the colors represent (arbitrary units). Corresponding heatmap with gene expression in each individual cell can be found in Supp. Fig. S2B. (F) Plot of percentage reads per cell that map to each of three types of KSHV genes (latent, early and late) in each cluster in the lytic + vehicle sample. Clusters are sorted based on the total average percentage of KSHV reads. (G) Proportion of cells that map to each cluster in the four samples. (H) Heatmap of average expression of KSHV genes in each cluster in the lytic + IDN-6556 and lytic + IDN-6556 + anti-IFN samples. The legends on the right define what expression level the colors represent (arbitrary units). Corresponding heatmaps with gene expression in each individual cell can be found in Supp. Fig. 2B.

**Figure 4.**
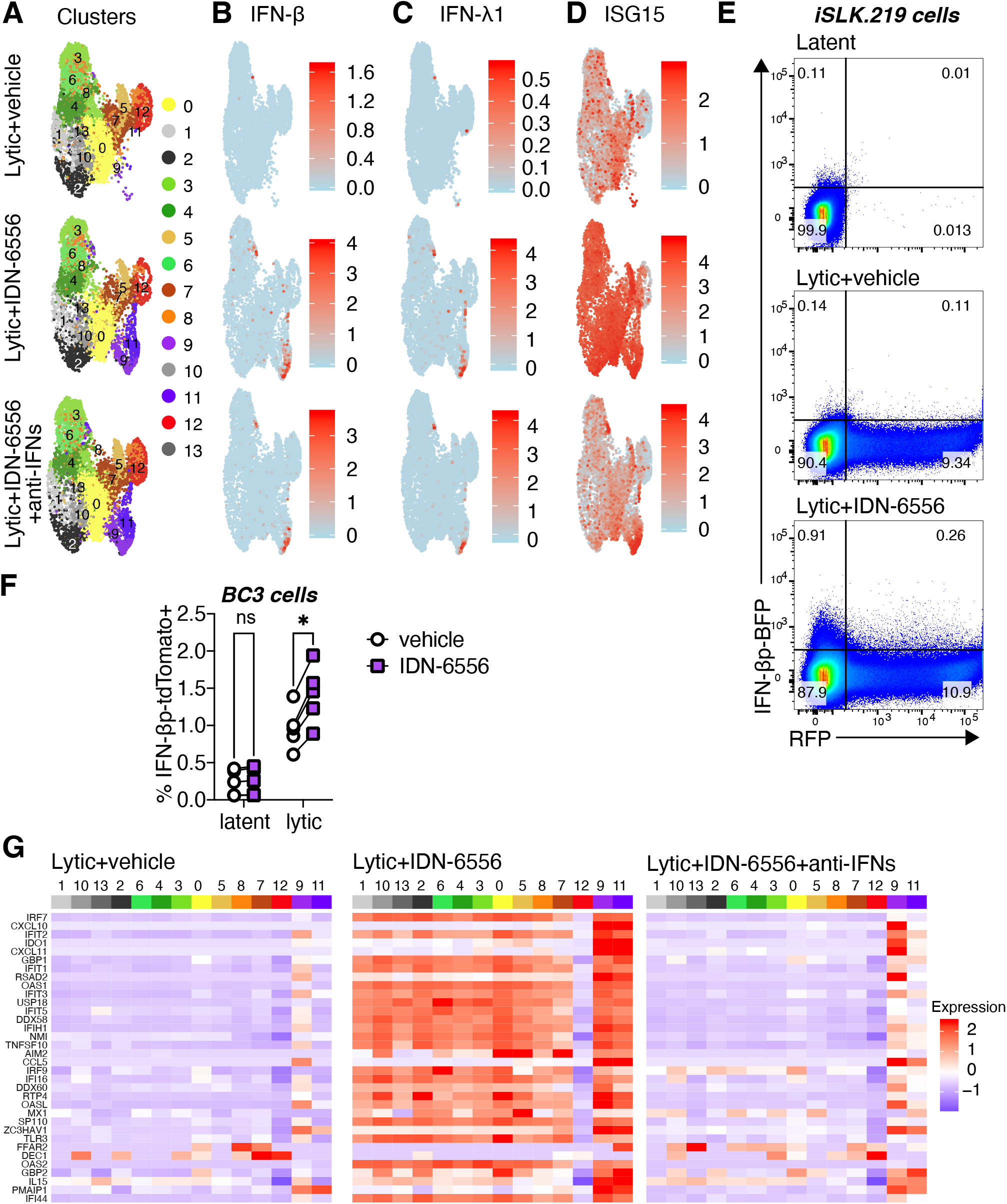
Only a small fraction of KSHV lytic cells express IFN-β. (A-D) Analysis of data from the scRNA-Seq experiment presented in Fig. 3A. (A) UMAP diagram of the clusters unbiasedly defined by Seurat in the 3 lytic samples (same as Fig. 3B). (B-D) UMAP diagram of the level of expression of IFN-β (B), IFN-l1 (C) or ISG15 (D) in each cell for the 3 lytic samples. The legends on the right define what expression level the colors represent (arbitrary units). (E) iSLK.219 expressing TagBFP from a human IFN-β promoter were treated with doxycycline to reactivate the lytic cycle (“lytic”) and IDN-6556, where indicated. Flow cytometry was carried out on the cells 4 days after doxycycline addition. RFP axis reflects the RFP signal in lytic cells and BFP axis reflects activity of the IFN-β promoter. Percentage of cells mapping to each quadrant are listed. Flow cytometry plots are representative of 3 repeats. (F) BC3 expressing tdTomato from a human IFN-β promoter were treated with TPA to reactivate the lytic cycle (“lytic”) and the caspase inhibitor IDN-6556, where indicated. Flow cytometry was carried out on the cells 2 days after TPA addition. The percentage of cells that were tdTomato positive is plotted for 5 biological replicates. ^ns^, **: *p* value > 0.05, < 0.01, Šidák’s multiple comparisons test after two-way ANOVA. (G) Analysis of data from the scRNA-Seq experiment presented in Fig. 3A. Heat maps of average expression of ISGs in each cluster for the three lytic samples. The legends on the right define what expression level the colors represent (arbitrary units). Corresponding heatmaps with gene expression in each individual cell can be found in Supp. Fig. 5B.

### Only a small subset of infected cells expresses IFNs to confer a potent antiviral state

An outstanding question from our bulk analysis was the identity of the cells that produce IFN-β in the caspase inhibitor-treated samples. We had hypothesized that lytically reactivating cells, rather than latent ones, produce IFNs. Indeed, our scRNA-Seq data showed that type I IFN expression is limited to cells that express viral lytic genes (Fig. 3H). However, the scRNA-Seq also surprisingly revealed that only 3.8% of the cells in the caspase inhibitor-treated samples expressed IFNs, particularly IFN-β (Fig. 4B). The IFN-β+ cells were concentrated in clusters 9 and 11, which only appeared in the +IDN-6556 samples (Fig. 4A, B). We did not detect expression of IFN-αs, but we saw that cells in clusters 9 and 11 also expressed the type III IFN IFN-λ1 (Fig. 4C, Supp Fig 3A). The IFN coming from this small fraction of cells was sufficient to elicit a strong IFN response in the entire population of cells, as indicated by the IFN-dependent increase in expression of IFN-stimulated genes (ISGs) after IDN-6556 treatment across almost all clusters (Fig. 4D, E, Supp. Fig. 5B). The only exception was cluster 12, which contains cells expressing late lytic gene. These cells are likely undergoing host shutoff, and generally express reduced levels of host genes(26). To ensure that the variability in IFN-β induction is not an artifact of a limit of detection by scRNA-Seq, we constructed iSLK.219 and BC3-based cell lines that expressed the TagBFP or tdTomato proteins under the control of the IFN-β promoter. When the lytic cycle was induced in these cells concomitantly with caspase inhibitor treatment, we found that ∼1-2% of the cells became TagBFP+ (Fig. 4E) or tdTomato+, in line with our scRNA-Seq results (Fig. 4F). In addition, we found that caspase inhibitor treatment of latent BC3 cell lines did not increase the percentage of tdTomato+ cells (Fig. 4F), consistent with our previous results that lytic induction is also required for the type I IFN induction (Fig. 2B, 2D, (4)). Overall, these data show that an exceedingly small fraction of cells induce IFNs even under caspase inhibition, yet they can elicit a strong anti-viral response in the entire cell population. Moreover, this heterogeneous induction of type I IFN occurs in both cell types we tested.

### The heterogeneity of type I IFN induction is not easily explained by differential viral gene expression

We considered the possibility that only a subset of KSHV-infected cells makes IFN-β due to differential expression of viral genes. For example, IFN-β+ cells could lack expression of KSHV-encoded anti-IFN factors. Consistent with this idea, cells in the IFN-β+ clusters (clusters 9 and 11) expressed only early lytic genes and low levels of the RFP lytic marker (Fig. 3H, 5A, Supp. Fig. 2B, 5C). However, their viral gene expression pattern was similar to that of other clusters that do not express IFNs, especially cluster 0 (Fig. 3H, 5A, Supp. Fig. 2B, 5C). Moreover, the viral gene expression pattern is also similar between IFN-β- and IFN-β+ cells within cluster 9 and 11 (Fig. 5A, Supp. Fig. 5C). One notable exception was the K5 mRNA, which was differentially expressed depending on IFN status. However, expression of K5 was actually higher in the IFN-β+ cells, rather than lacking in these cells. These observations suggest that lack of expression of viral anti-IFN regulators does not explain the restricted IFN-β induction. Another possibility is that some cells express higher basal levels of genes in the type I IFN induction pathway and are thus more sensitive to IFN-inducing cues. This model was proposed in the case of other viral infections by Zhao *et al*.(27). Analyzing these genes is complicated by the fact that many of them are themselves ISGs. Indeed, their expression was increased in all clusters (except cluster 12) in the caspase inhibitor-treated samples by IFN signaling, and was reduced back to basal levels by anti-IFN antibody treatment (Fig. 5B, Supp. Fig. 5B). Nonetheless, there was a clear enrichment of transcripts for the NF-κB transcription factor family members (NFKB1, NFKB2, RELA, RELB) in the IFN-β+ clusters 9 and 11 (Fig. 5B, Supp. Fig. 5B). NF-κB is also needed for IFN-β transcription downstream of pathogen sensors, together with IRF3(1). Indeed, levels and phosphorylation of NF-κB were increased in lytic cells treated with IDN-6556, showing that caspase activity also modulates NF-κB activation (Fig. 5C). It is thus possible that high NF-κB expression is needed to induce type I IFN in these cells, perhaps as a feed-forward mechanism to ensure commitment to IFN production.

**Figure 5.**
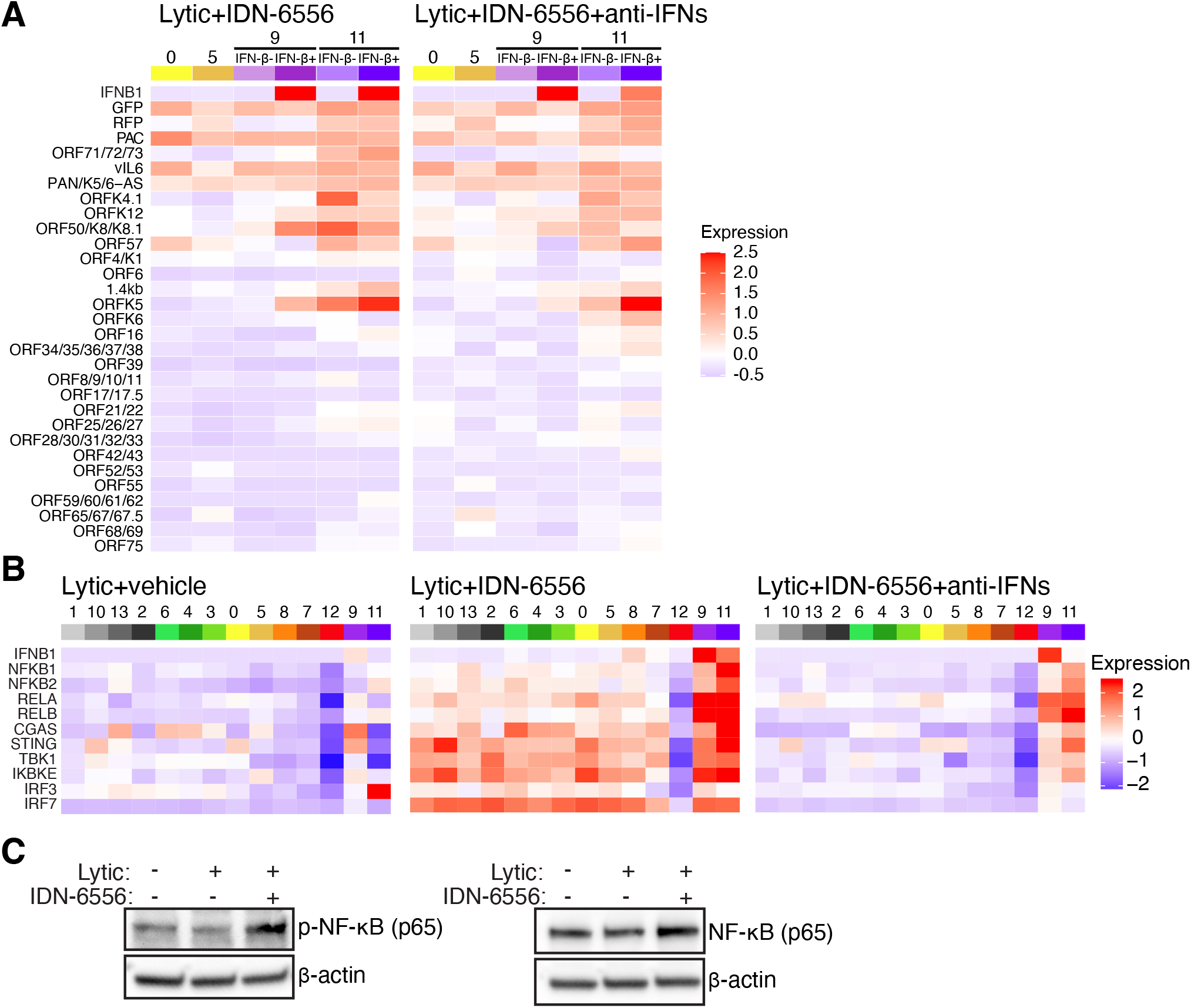
IFN-β-producing cells express high levels of NF-κB transcripts. (A-B) Analysis of data from the scRNA-Seq experiment presented in Fig. 3A. Heatmaps of average expression of KSHV genes in clusters 0, 5, 9, and 11, with 9 and 11 divided based on IFN-β status (A) and of average expression of select genes from the type I IFN induction pathway (B) in each cluster for the three lytic samples. The legends on the right define what expression level the colors represent (arbitrary units). Corresponding heatmaps with gene expression in each individual cell can be found in Supp. Fig. 5B and 5C. (C) iSLK.219 were treated with doxycycline to reactivate the lytic cycle (“lytic”) and with IDN-6556 where indicated. Protein lysates were collected at day 4 and analyzed by western blotting for NF-κB, phospho-NF-κB (p-NF-κB) and β-tubulin as a loading control. Blots are representative of 3 replicates.

## Discussion

This study provides mechanistic insights into multiple levels of type I IFN regulation during KSHV infection. We find that caspases regulate cGAS activity during KSHV lytic infection, likely indirectly, either through a cGAS regulator/cofactor or by modulating access of cGAS to its stimulating DNA (Fig. 1-2). Moreover, we find that although caspase inhibition results in a strong type I IFN induction, type I and III IFNs are only turned on in a very small percentage of cells. IFN expression appears to correlate with strong induction of NF-κB transcripts, rather than differences in viral gene levels.

The cGAS/STING pathway has emerged in recent years as a crucial component of the innate immune response to DNA viruses and various other pathogens(28). This is highlighted by the multitude of strategies employed by pathogens(29), including herpesviruses(30), to disrupt the cGAS/STING pathway. While this immunosuppression is often driven by virus-encoded proteins, we have uncovered an intriguing mechanism in KSHV infection, which involves the hijacking of the host cell caspases to subvert immune activation and viral clearance(4). During apoptosis and other viral infection, caspase-3 and -7 have been reported to directly cleave cGAS (8). However, our previous results suggest that caspase-8 activity mediates type I IFN suppression in KSHV-infected cells without triggering apoptosis, whereas caspase-3/7 activity is dispensable(4). Moreover, we have been consistently unable to detect cleavage fragments or downregulation of cGAS in infected cells. Together, our previous report and the current study suggest an unexpected type I IFN regulatory function of caspase-8 activity in targeting the cGAS/STING pathway that is separate from its apoptotic activity. Perhaps caspases like caspase-8 cleave regulators of the cGAS/STING pathway. Alternatively, they may affect access of the DNA to cGAS. The latter is hard to test, because we do not know what the cGAS stimulating DNA is during KSHV reactivation. We have previously reported that viral DNA replication is not required for IFN induction (4). Moreover, we have found that depletion of mitochondrial DNA, a common stimulus of cGAS, reduced reactivation of the lytic cycle, confounding analysis of IFN induction (data not shown). Identifying potential relevant caspase targets will be important to determine how they regulate cGAS activity. Nonetheless, caspases are an attractive therapeutic target that could be exploited to potentiate cGAS/STING-mediated immune responses for viral clearance. This could have far reaching implications for other pathologies, as regulation of the cGAS/STING pathway is fundamental in various inflammatory diseases and in tumor immunity(31, 32), and thus is a current focus of intensive drug discovery research efforts.

The notable cellular heterogeneity of type I IFN production reveals an additional layer of complexity and regulation of innate immune responses during KSHV infection. We detect an exceptionally small fraction of IFN-β+ cells upon caspase inhibitor treatment (< 5%, Fig. 4B). Remarkably, this small group of IFN-β+ cells still induces a robust IFN response in the whole population (Fig. 4D, F). On a practical level, the high heterogeneity in IFN-β expression complicates our analysis of caspase regulation of cGAS. It is possible that the upstream portions of the pathway (i.e. cGAS) are also only active in the IFN-β+ cells, and that cell sorting may be needed to detect differences in the levels of the direct target(s) of caspases in IFN regulation. Alternatively, it is possible that only a subset of the cells in which the IFN induction pathway is activated end up expressing IFN-β. In support of this idea, we were able to easily detect phosphorylation of many of the components of the IFN induction pathway using bulk analysis (Figs. 1E, 2C, 5C).

Single-cell RNA-Seq studies in herpesviruses have started to come out in the last few years, but many have not analyzed IFN responses, focusing instead on latency and reactivation (33–38). However, several studies in multiple viruses have examined IFN induction, and have also showed that only a small number of cells respond to infection in a population during infection with a number of other viruses (27, 35, 39–45). Interestingly, studies that investigated infections in innate immune cells like macrophages and dendritic cells or fibroblasts saw higher percentages of responding cells (10-30%) (27, 39, 40). In contrast, iSLK.219 cells are of epithelial origin(46). Epithelial cells are often the first exposed to invading pathogens and mount responses that alert and prime surrounding cells to counteract viral spread. Therefore, it is possible that epithelial cell responses are even more tightly regulated to prevent inadvertent activation of the pathway. Consistent with this idea, influenza A virus and SARS coronavirus 2 infection of epithelial cells also elicit responses in a small minority of cells (41–43). Moreover, while we did not perform scRNA-Seq on BC3 cells, flow cytometry analysis of BC3 cells encoding an IFN reporter also indicate that IFN-β is induced in less than 2% of these cells (Fig 4F). Taken together, our findings suggest differences in IFN responsiveness depending on the cell type as well as the virus infecting the cells.

It is unclear what the source of interferon heterogeneity is in KSHV-infected cells. Viral gene expression, particularly expression of viral immune regulators, is often considered the reason for limited type I IFN responses(43). A previous study in herpes simplex virus 1 reports that type I IFN signaling only occurs in a small percentage of cells that are all abortively infected(35). This is also a potential explanation for our results, since we find that the cells that make IFNs only express a subset of viral genes, mostly early genes (Fig. 3H, 5A). However, this pattern of gene expression is not unique to the small percentage of IFN-β+ cells. Multiple clusters of cells do not express high levels of late lytic genes. Indeed, the pattern of viral gene expression is generally similar in clusters 0, 5 (IFN-β negative) and 9, 11 (IFN-β positive) (Fig. 5A, Supp. Fig. 5C). Moreover, when we compare IFN-β positive and negative cells within the same clusters (clusters 9 and 11), we find that they also have a similar viral gene expression pattern (Fig. 5A, Supp. Fig. 5C). We also do not know whether this gene expression pattern represents abortive infection, or simply an early stage of reactivation that facilitates IFN induction, since we know that iSLK.219 reactivation is heterogeneous. This also means that we cannot distinguish a situation whereby progression to the late stage of the lytic cycle prevents IFN-β induction from one where IFN-β induction prevents progression through the lytic cycle. It thus appears that abortive or early stage infection alone does not explain the restricted pattern of IFN gene expression in our experiments. Our results also do not support the idea that some cells express IFN because they are missing expression of a viral negative regulator of type I IFN, although we cannot fully exclude this model because we do not reliably detect all viral genes. However, we did find that the KSHV gene K5 displayed much higher expression in the IFN-β-positive cells (Fig. 5A, Supp. Fig. 5C). This was surprising to us as we would not expect viral genes to exhibit positive regulation on IFN induction. There is no known role of K5 in regulating type I IFN signaling, so it is unclear why its expression correlates with IFN-β expression at present. Future studies will hopefully elucidate this correlation.

Currently, our results point to the possibility that inherent differences in the cells govern the heterogeneity of the interferon response. Stochasticity in gene expression, which is common among cytokines, has previously been proposed as driver of type I IFN expression heterogeneity. Zhao *et al*. suggested some of this stochasticity may be epigenetically encoded(27), reducing the number of cells that are able to express type I IFN, perhaps to prevent over-activation and inflammation. Paracrine IFN signaling is also thought to influence and modulate heterogeneity(44). This cell-cell communication may fine tune immune signal amplification in a cell density-dependent manner. Indeed, we observe 3.8% IFN-β+ cells in the IDN-6556 treated sample, but only 1.6% IFN-β+ cells in the sample treated with IDN-6556 and anti-IFN antibodies that block IFN signaling. Moreover, while cells in cluster 9 retain high average IFN-β expression even when IFN signaling is blocked, cluster 11 cells have lower IFN-β without IFN signaling (Fig. 5A, Supp. Fig. 5A, C), suggesting two different modes of IFN induction in these groups of cells. This reinforces the notion of self-amplification of the type I IFN response as a mechanism to ensure robust responses only in the appropriate contexts for infection control, while avoiding tissue damage from unnecessary inflammation.

## Materials and Methods

### Cell lines, constructs, reagents, and treatments

All cells were cultured at 37 °C and 5% CO_2_ conditions. iSLK.219, iSLK.219-IFNBp-BFP, iSLK.RTA(16), and HEK293T cells were grown in Dulbecco’s modified Eagle’s medium (DMEM; Life Technologies) supplemented with 10% fetal bovine serum (FBS) (HyClone). BC3 cells were grown in RPMI supplemented with 20% FBS, 2 mM GlutaMAX supplement, and 50 μM β-mercaptoethanol (Gibco/Thermo Fisher). iSLK.219 cells were reactivated with 1 μg/ml of doxycycline (Thermo Fisher). Reactivation was confirmed visually for all iSLK.219 samples by examining the appearance of RFP-positive cells (RFP is driven by the lytic PAN promoter). BC3 cells were reactivated with 20 ng/ml of TPA for 48 h. iSLK.219-IFN-βp-BFP cells were generated by transducing iSLK.219 cells with lentiviruses containing pJP1_IFNBp-tagBFP and BC3-IFN-βp-tdTomato by transducing BC3 cells with pLJM1_2xIFNBp-tdTomato. All pLJM and pJP1 are based on pLJM1-GFP, a gift from David Sabatini (Addgene plasmid # 19319 ; http://n2t.net/addgene:19319 ; RRID:Addgene_19319)(47). For pJP1_IFNBp-tagBFP, the CMV promoter of pLJM1 was replaced with a human IFN-β promoter fragment subcloned from a luciferase reporter construct (kind gift of Michaela Gack) and the GFP sequence was replaced with a TagBFP sequence. In addition, the antibiotic resistance gene was changed from a puromycin resistance to a zeocin resistance gene. For pLJM1_2xIFNBp-tdTomato, the CMV promoter and multiple cloning sites were replaced with two copies of a human IFN-β promoter 303 bp fragment cloned from IFN-Beta_pGL3, a gift from Nicolas Manel (Addgene plasmid # 102597 ; http://n2t.net/addgene:102597 ; RRID:Addgene_102597)(48). To add the 54 bp 5’ untranslated region of human IFN-β, sequences were added to the primers used to assemble the construct. Lentiviral packaging was carried out using packaging plasmids pMDLg/pRRE (Addgene plasmid # 12251 ; http://n2t.net/addgene:12251 ; RRID:Addgene_12251), Addgene plasmid # 12253 ; http://n2t.net/addgene:12253 ; RRID:Addgene_12253) and pMD2.G (Addgene plasmid # 12259 ; http://n2t.net/addgene:12259 ; RRID:Addgene_12259), kind gifts from Didier Trono(49). Where indicated, cells were treated with vehicle (dimethyl sulfoxide [DMSO]; Sigma-Aldrich), 10 μM IDN-6556 (MedChem Express or Selleck Chemicals), 24.1 µM RU.521 (Invivogen), 1:50, 1:100, or 1:500 dilutions of a mixture of neutralizing antibodies against type I IFNs (PBL Assay Science, catalog # 39000-1). The concentrations of all drug treatments were chosen based on common concentrations used in previously published studies.

### siRNA knockdown

iSLK.219 cells were transfected while in suspension (reverse transfection) with 10 nM siRNAs (purchased from Thermo Fisher) against STING (HSS139156), MAVS (HSS148537), cGAS (siRNA#1, HSS132955; siRNA#2, HSS132956), IFI16 (siRNA#1, HSS105205; siRNA#2, HSS105206), RIG-I (siRNA #1, s24144; siRNA #2, s223615), or a nontargeting siRNA (12935300) at a density of 16,700 cells/cm^2^ using the Lipofectamine RNAiMAX transfection reagent (Life Technologies/Thermo Fisher) per the manufacturer’s protocol. Six hours later, the medium was replaced. The lytic cycle was induced two days later. For cGAS and RIG-I knockdowns, the reverse transfection process was done twice, the first time two days prior to induction and a second time on the day of lytic cycle induction. All siRNAs were obtained from Life Technologies/Thermo Fisher.

### Realtime quantitative polymerase chain reaction (RT-qPCR)

For RNA analysis, iSLK.219 cells were seeded at a density of 33,000 cells per well in a 24-well plate, and the lytic cycle was induced by the addition of doxycycline (1 μg/ml). RNA samples were collected at the indicated time points in RNA lysis buffer (Zymo Research). BC3 cells were seeded at 500,000 cells per ml and induced using TPA for 48 h. One ml of the cells was collected, and the cell pellet was lysed in RNA lysis buffer (Zymo Research). Total RNA was extracted using the Quick-RNA MiniPrep kit (Zymo Research) by following the manufacturer’s protocol. For mRNA measurements, cDNA was prepared using an iScript cDNA synthesis kit (Bio-Rad) per the manufacturer’s protocol. In all cases, 18S rRNA levels were used as an internal standard to calculate relative mRNA levels. Real-time quantitative PCR (RT-qPCR) was performed using iTaq Universal SYBR Green Supermix (Bio-Rad) in a CFX Connect real-time PCR detection system (Bio-Rad). No-template and no-RT controls were included in each replicate. CFX Manager software was used to analyze the data. Primers are listed in Table S3.

### Protein analysis

iSLK.219 cells were seeded at a density of 167,000 cells per well in a 6-well plate and treated with doxycycline (1 μg/ml). Lysates were collected at the indicated time points. Cells were lysed in radioimmunoprecipitation assay (RIPA) buffer (50 mM Tris-HCl, pH 7.4, 150 mM NaCl, 2 mM EDTA, 0.5% sodium deoxycholate, 0.1% SDS, 1% NP-40) or an NP-40-only buffer for caspase blots and phospho-protein blots (50 mM Tris-HCl, pH 7.4, 150 mM NaCl, 1 mM EDTA, 0.5% NP-40) supplemented with 50 μg/ml phenylmethylsulfonyl fluoride (PMSF; G-Biosciences) and cOmplete protease cocktail inhibitor (Roche). Samples were separated by SDS-PAGE and transferred to polyvinylidene difluoride (PVDF) membranes. The following Cell Signaling Technologies antibodies were used on PVDF membranes at 1:1,000 dilution in 5% milk in in Tris-buffered saline with 0.1% Tween 20 (TBST): anti-cGAS (D1D3G) (no. 15102). The following Cell Signaling Technologies antibodies were used with PVDF at 1:1,000 dilution in 5% BSA in TBST: anti-caspase-8 (no. 9746), anti-cleaved caspase-8 (no. 9496), anti-IRF3 (D83B9) (no. 4302), anti-phospho-IRF3 Ser396 (4D4G) (no. 4947), anti-TBK1 (D1B4) (no. 3504), anti-phospho-TBK1 Ser172 (D52C2) (no. 5483), anti-STING (D2P2F) (no. 13647), anti-phospho-STING Ser366 (D7C3S) (no. 19781), anti-RIG-I (D14G6) (no. 3743), anti-NF-κB p65/RelA (D14E12) (no. 8242S), anti-phospho-NF-κB p65 Ser536/RelA (93H1) (no. 3033) and anti-β-tubulin (9F3) (no. 2128). Anti-IFI16 (ab169788; 1:1000, Abcam) and anti-β-actin (ab8229; 1:500, Abcam) were diluted in 5% BSA in TBST and used with PVDF membranes blocked in 5% BSA in TBST. Anti-MAVS (sc-166583, 1:1000 dilution; Santa Cruz Biotechnology) was diluted in 0.5% milk in phosphate-buffered saline with 0.1% Tween 20 (PBST) and used with PVDF membranes blocked in 5% milk in PBST. Horseradish peroxidase (HRP)-conjugated goat anti-rabbit IgG and goat anti-mouse IgG (both 1:5,000 in blocking buffer) were purchased from Southern Biotechnology. HRP-conjugated donkey anti-goat IgG (1:5,000 in 0.5% milk in PBST) was purchased from Santa Cruz Biotechnology. All membranes were exposed using Pierce ECL Western blotting substrate (Thermo Fisher) and imaged with a Syngene G:Box Chemi XT4 gel doc system.

### cGAMP ELISA

Cell lysates were harvested 4 days post reactivation in RIPA and subjected to enzyme-linked immunosorbent assay (ELISA) analysis for 2’3’-cGAMP (Cayman Chemical) according to the manufacturers’ protocols. Each biological replicate consisted of three technical replicates per condition.

### Flow cytometry analysis

10^6^ IFN-βp-TagBFP cells were treated with DMSO, IDN-6556 (10 μM), docycline (1 mg/ml) or doxycycline and IDN-6556 for 4 days to induce KSHV reactivation. 10^6^ BC3-IFN-βp-tdTomato cells were treated with DMSO, IDN-6556 (10 μM), TPA (20 ng/mL), or TPA and

IDN-6556 for 48 hours to induce KSHV reactivation. Cells were fixed in 4% PFA for 15 minutes at room temperature and washed in PBS before being analyzed on a BD Biosciences LSR II flow cytometer in the Tufts Laser Cytometry Core facility. 500,000 events were collected per sample. TagBFP, RFP, and tdTomato expression was gated based on the corresponding latently infected and vehicle (DMSO) treated cells.

### Single cell RNA sequencing

iSLK.219 and iSLK.RTA cells were seeded onto T25 flasks at a density of 16,700 cells/cm^2^ and induced with doxycycline in the presence or absence of IDN-6556 (10 μM) and a mixture of neutralizing antibodies against type I IFNs at a 1:500 dilution (39000-1; PBL Assay Science). The media was replaced at day 2 post reactivation with fresh doxycycline, IDN-6556, and type I IFN neutralizing antibodies. Cells were harvested at day 4 post reactivation, washed twice with PBS, and strained to create single cell suspensions at a concentration of 1,000,000/ml. Single cell samples were generated according to 10X Genomics’ sample preparation protocol for cultured cell lines (CG00054) and Gel Beads-in-emulsion (GEM) generation protocol (CG000183). Briefly, cells were trypsinized and washed twice with Dulbecco’s phosphate buffer saline (DPBS, Life Technologies). Cells were resuspended in DPBS and passed through a 30 µm strainer to remove clumps. Cells were counted and diluted to a concentration of 1,000,000 cells/mL. Cells were mixed with a reverse transcription mastermix (including a primer with polydT anchor) and loaded onto a Chromium B microfluidic chip. Gel beads and partitioning oil were loaded onto the chip and the chip was run through the chromium controller device to generate the GEMs. GEMs were incubated in a thermal cycler for 1 cycle to generate cDNA. The emulsion was broken, and cDNA was isolated with magnetic beads, amplified for 12 cycles, and then cleaned up with SPRIselect reagent.

Libraries were prepared according to 10X Genomics library construction protocol (CG000183). Briefly, cDNA was fragmented and then end-repaired and A-tailed with the enzymes in the provided 10X Genomics kit (version 3, PN-1000092). cDNA was size selected using SPRIselect and adaptor oligos were ligated onto the cDNA and further cleaned up with SPRIselect. Paired-end sequencing with 75 cycle read length was performed on an Illumina Nextseq 550 at Tufts Genomics Facility. Raw sequencing data (.bcl files) were processed using the 10X Genomics CellRanger (3.0.0) software. Fastq files were generated for the reads using the mkfastq function as described on the 10X Genomics website (https://support.10xgenomics.com/single-cell-gene-expression/software/pipelines/latest/using/mkfastq). Also using CellRanger, the count function was used to align the reads to a reference genome including the human genome (GRCh38), the KSHV genome (GQ994935.1), and the additional sequence of the RFP/GFP/PAC insertion cassette (Genbank MZ617352). This third reference was added because the GQ994935.1 reference genome corresponds to the KSHV BAC16 genome, whereas our cells contain the rKSHV.219 virus(16, 50). To our knowledge, a full reference for rKSHV.219 is not available. We thus amplified the missing region and sequenced it using Sanger sequencing. The matrix table of expression for each gene in each single cell from CellRanger was used as an input for integration and clustering analyses using the R-based Seurat package (version 3.1.1.9021, R version 3.6.1)(19, 20). Cell cycle analysis was done according Tirosh *et al*.(51). Raw and processed data are available on NCBI GEO, record # GSE190558. Scripts used for analysis are available on GitHub, https://github.com/mgaglia81/iSLK.219_scRNAseq. The matrices were converted to Seurat objects and then pre-processed by performing quality control analyses. First, the four objects from the different samples were integrated together into one object using Seurat functions FindIntegrationAnchors() and IntegrateData(). Dying cells with mitochondrial contamination or possible cell doublets were filtered out by eliminating cells that have >20% mitochondrial RNA counts and cells that have unique feature (RNA) counts > 7500. Feature expression measurements for each cell were normalized by the total expression, multiplied by a scale factor (10,000) and log-transformed using the Seurat function NormalizeData(). The expression was scaled so that the mean expression across all cells is 0 and the variance is 1, using the Seurat function ScaleData(). For linear dimensional reduction, principal component analysis (PCA) was performed using the Seurat function RunPCA(). Only the PCs that account for the most variation were included, to overcome technical noise when clustering cells. These were chosen by visualizing each PC with Seurat functions JackStrawPlot() and ElbowPlot(), and determining that variance was negligible after 30 PCs. Therefore, UMAP cluster plots were created using dimensions 1-30.

### Statistical analysis

All statistical analysis was performed using GraphPad Prism version 8.4.2 or later (GraphPad Software, La Jolla California USA, www.graphpad.com). Statistical significance was determined by Student’s *t*-test or analysis of variance (ANOVA) followed by a *post hoc* multiple comparison test (Dunnett’s, Sidak’s or Tukey’s) when multiple comparisons were required. All data are plotted as mean ± standard deviation.

## Acknowledgements

We thank members of the Gaglia laboratory for suggestions and feedback on the project and the manuscript. We thank the Tufts Laser Cytometry and Genomics core staff for technical and conceptual assistance. We thank Drs. Gack, Trono, Sabatini, and Manel for sharing reagents. This work was supported by American Cancer Society grant 131320-RSG-17-189-01-MPC and NIH R01 R01CA268976 to MMG. TT was supported by the Tufts Collaborative Cancer Biology Award. RCL was supported by NIH training grant T32 GM007310.

---

**Supplemental Figure 1.**
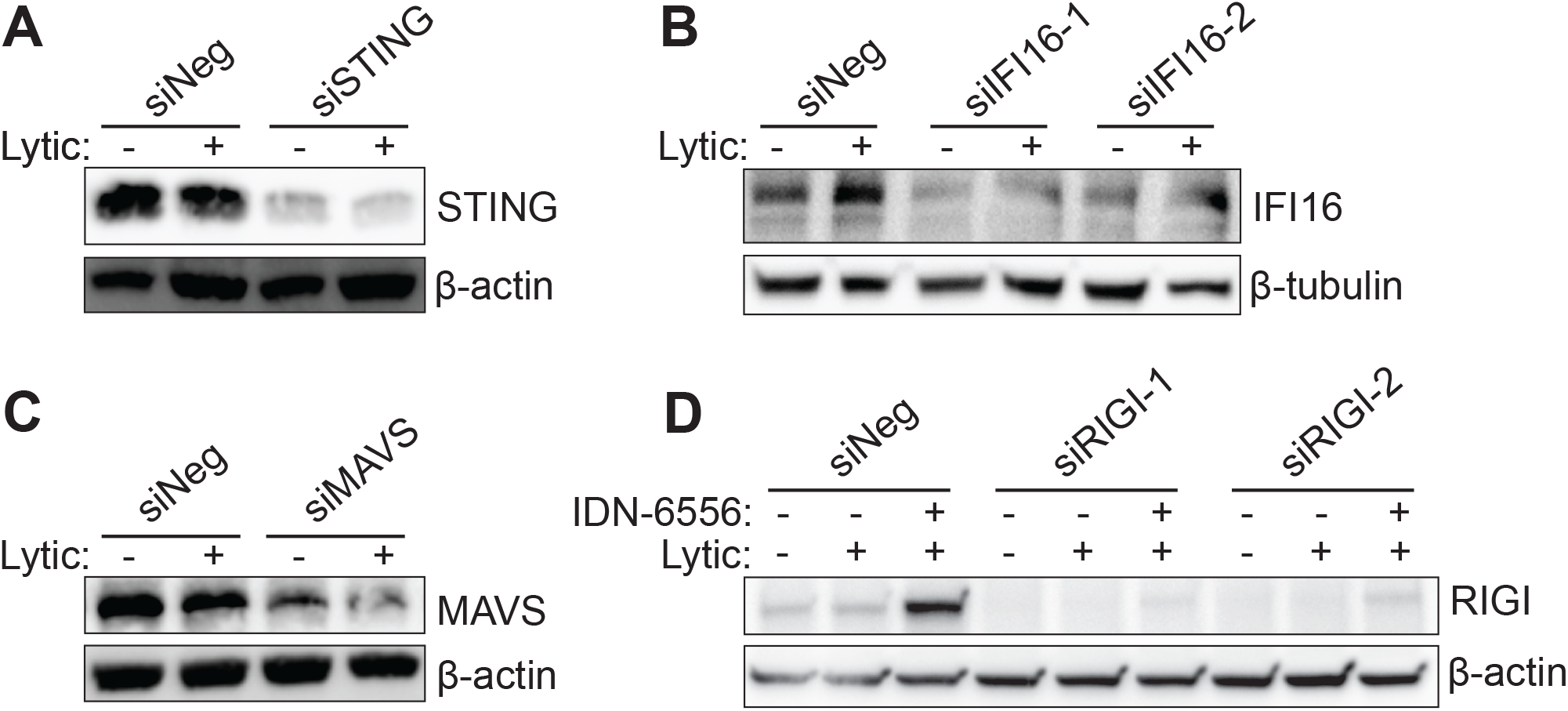
Knockdown of proteins in the type I IFN induction pathway. iSLK.219 cells were transfected with a negative control siRNA and siRNAs targeting STING (A), IFI16 (B), MAVS (C), or RIG-I (D). Cell lysates were harvested at day 4 post reactivation with doxycycline and treatment with IDN-6556 (where indicated) and subjected to western blotting for the target proteins and β-actin or β-tubulin as loading controls. Blots are representative of 3 replicates.

**Supplemental Figure 2.**
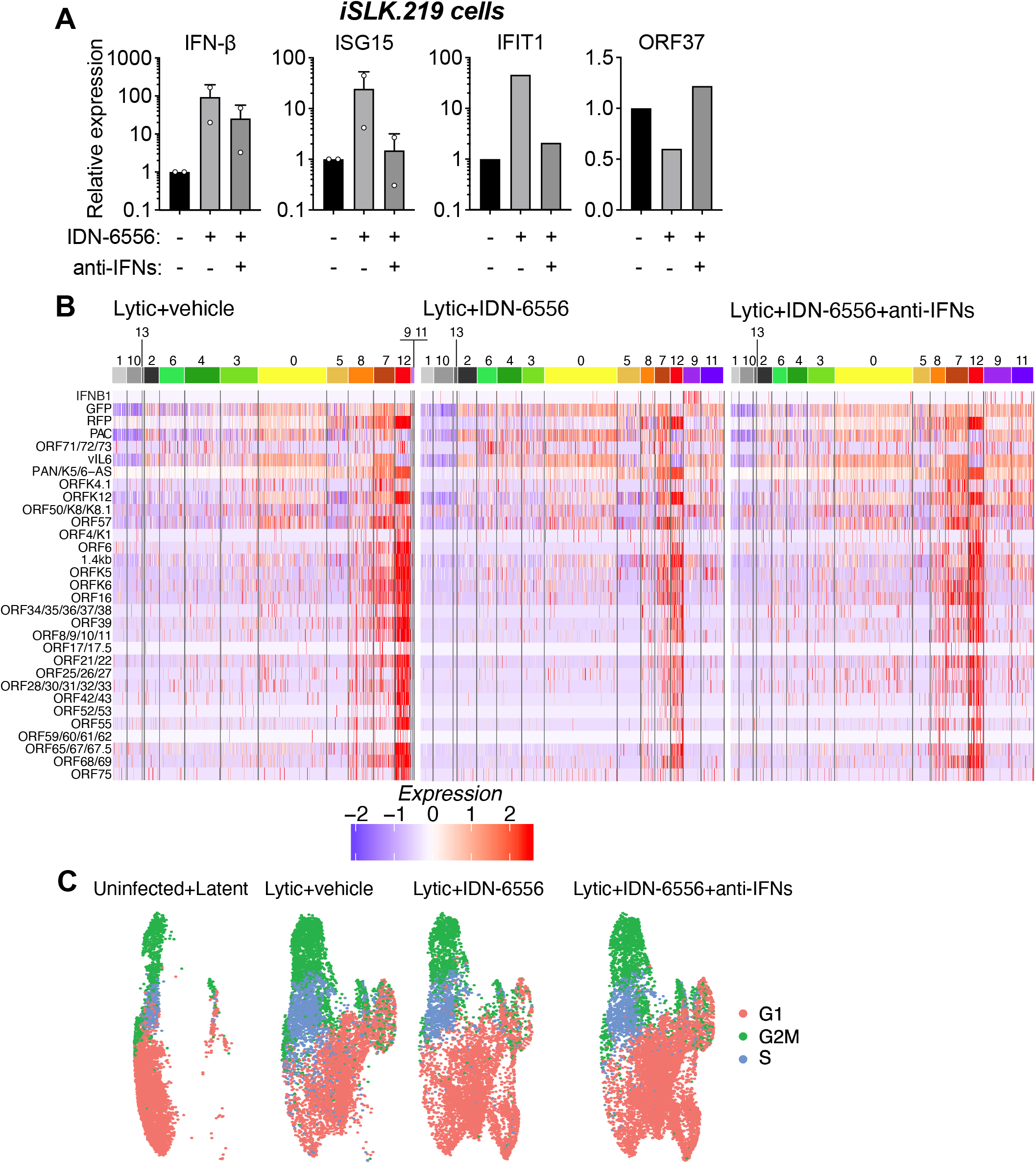
scRNAseq controls and additional analyses. (A) iSLK.219 cells were treated with doxycycline (1 μg/ml) to reactivate the lytic cycle, as well as IDN-6556 and anti-IFN antibodies where indicated. mRNAs levels of IFN-β, the ISGs ISG15 and IFIT1, and the KSHV gene ORF37 were measured by RT-qPCR four days after doxycycline addition. (B-C) Analysis of data from the scRNA-Seq experiment presented in Fig. 3A. (B) Heatmaps of expression of KSHV genes in each cell in the three lytic samples, sorted by cluster. The legend below the heatmaps defines what expression level the colors represent (arbitrary units). (G) UMAP diagram of cell cycle stage classification of cells in the 4 samples.

**Supplemental Figure 3.**
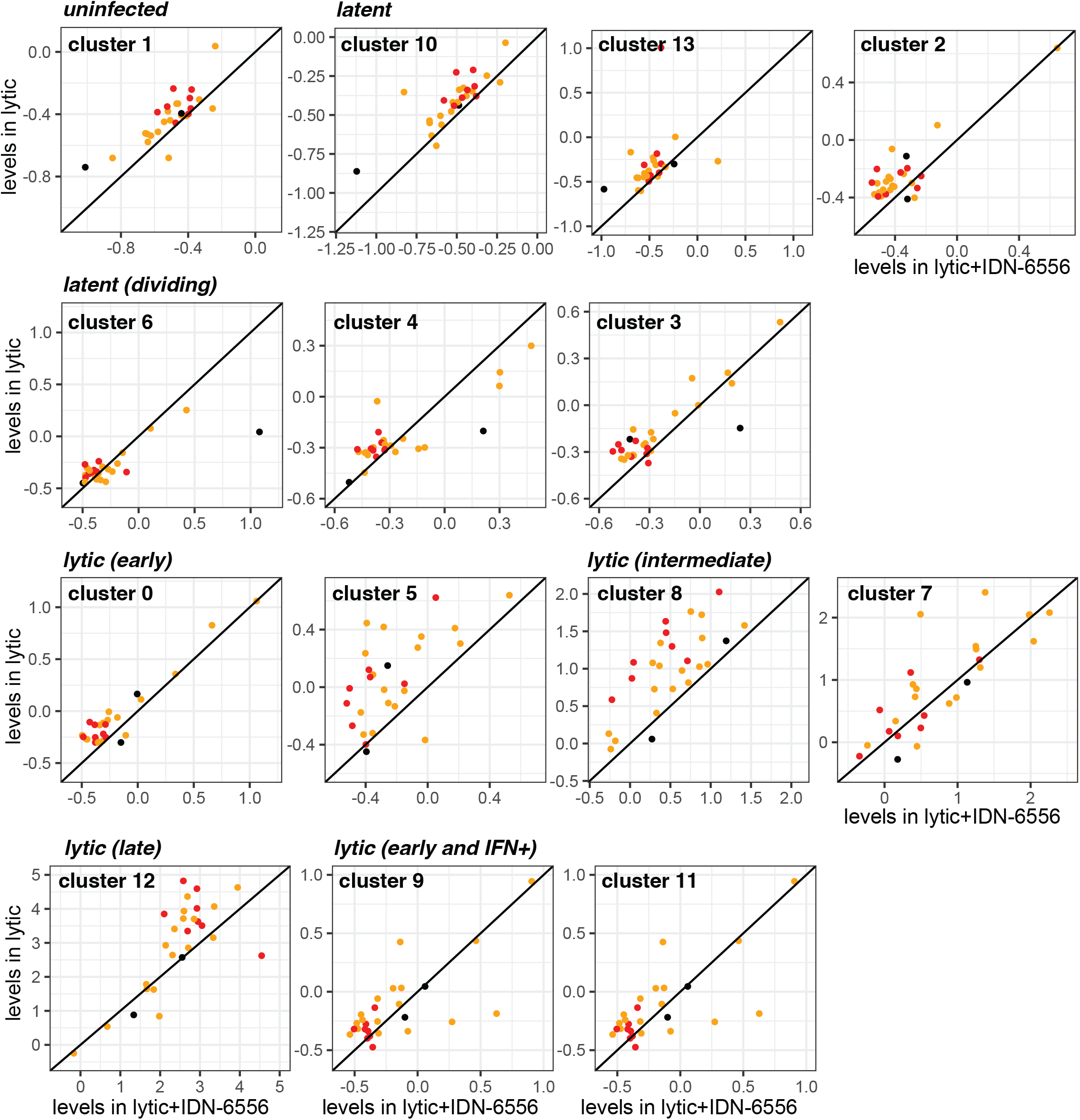
Changes in viral gene expression between lytic iSLK.219 cells and lytic iSLK.219 cells treated with caspase inhibitors. Analysis of data from the scRNA-Seq experiment presented in Fig. 3A. Scatter plots show the average expression of viral genes in each of the clusters (ordered based on our classification) in iSLK.219 cells treated with doxycycline (lytic, y-axis) and iSLK.219 cells treated with doxycycline and caspase inhibitors (lytic + IDN-6556, x-axis). Black dots represent latent genes, yellow dots early genes and red dots late genes. The diagonal line represents equal levels.

**Supplemental Figure 4.**
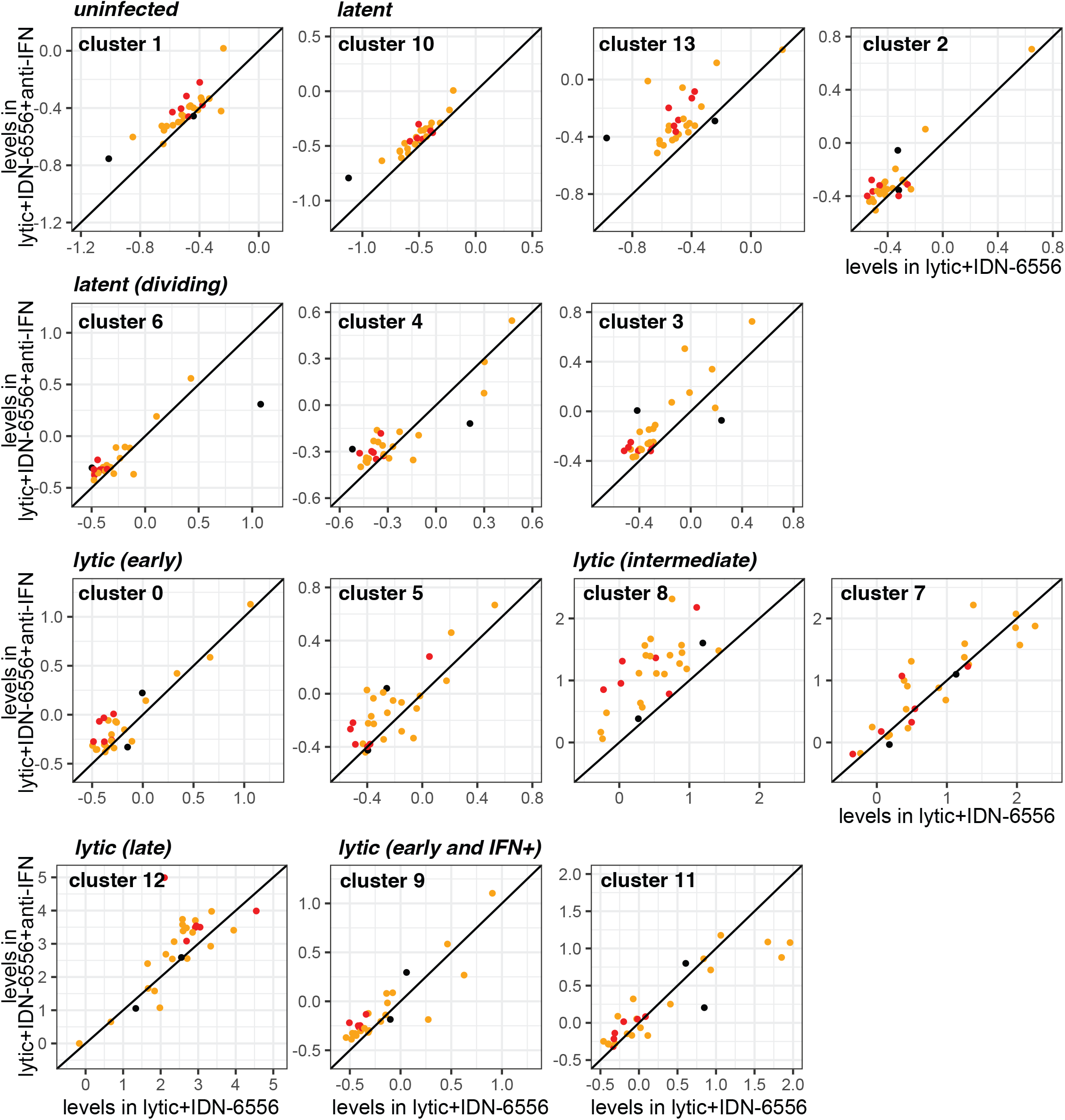
Changes in viral gene expression between lytic iSLK.219 cells treated with caspase-inhibitors and lytic iSLK.219 cells treated with caspase inhibitors and anti-IFN antibodies. Analysis of data from the scRNA-Seq experiment presented in Fig. 3A. Scatter plots show the average expression of viral genes in each of the clusters (ordered based on our classification) in iSLK.219 cells treated with doxycycline and caspase inhibitors (lytic + IDN-6556, x-axis) and iSLK.219 cells treated with doxycycline, caspase inhibitors, and anti-IFN antibodies (lytic + IDN-6556 + anti-IFN, y-axis). Black dots represent latent genes, yellow dots early genes and red dots late genes. The diagonal line represents equal levels.

**Supplemental Figure 5.**
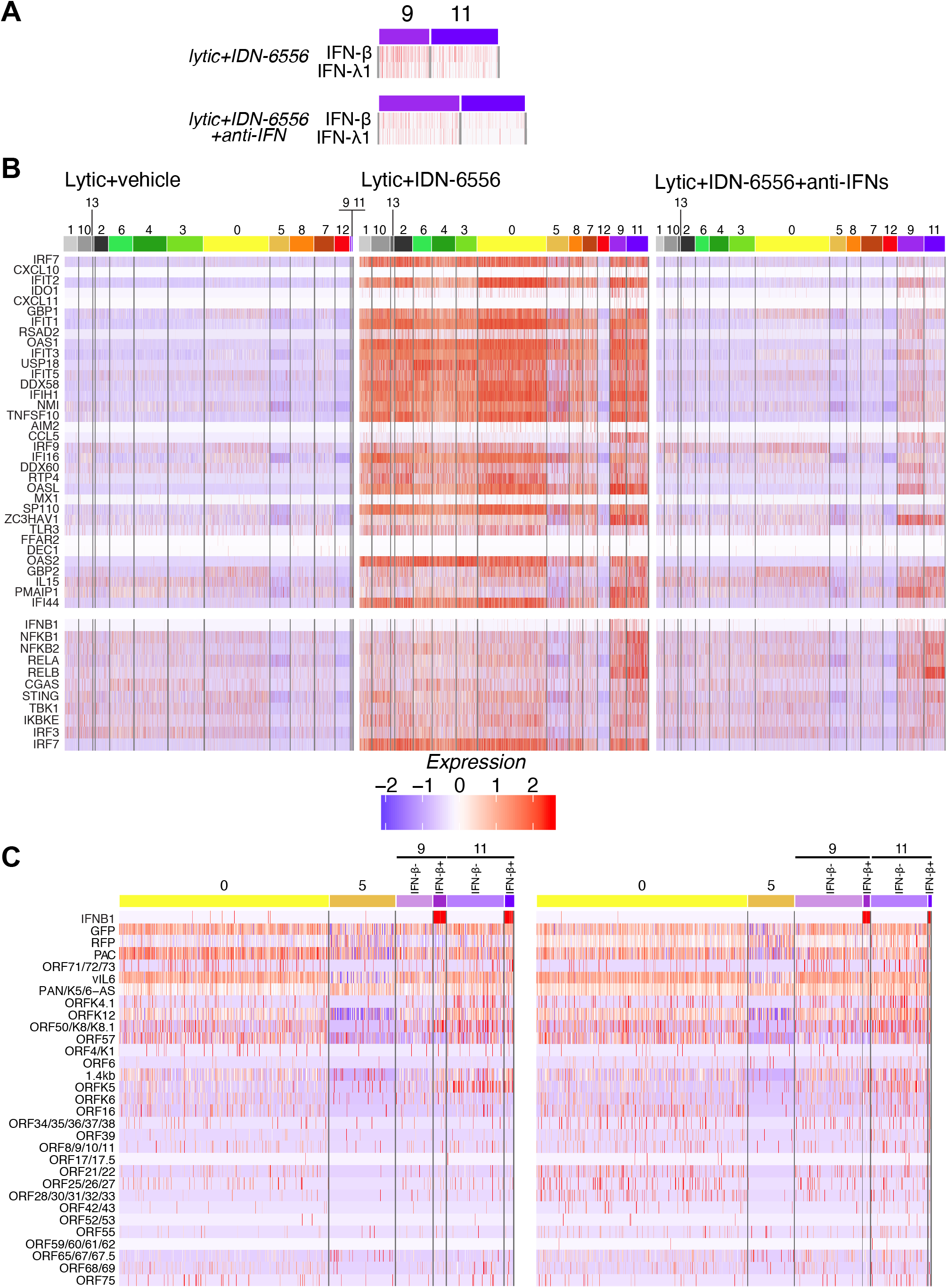
Expression patterns of IFNs, ISGs and genes in the type I IFN induction pathway. Analysis of data from the scRNA-Seq experiment presented in Fig. 3A. (A) Heatmaps of expression of IFN-b and IFN-l1 in each cell in clusters 9 and 11 in the two IDN-6556 treated samples. (B) Heatmaps of expression of ISGs (top) and select genes from the type I IFN induction pathway (bottom) in each cell in the three lytic samples, sorted by cluster. (C) Heatmaps of viral gene expression in each cell in clusters 0, 5, 9, and 11, with cells in 9 and 11 divided by IFN-b status, in the three lytic samples. The legend defines what expression level the colors represent (arbitrary units).

**Table S1.** Characteristics of scRNA-Seq datasets.

|  | Uninfected + latent | Lytic + vehicle | Lytic + IDN-6556 | Lytic + IDN-6556 + anti-IFNS |
| --- | --- | --- | --- | --- |
| Total reads | 142,704,565 | 138,925,899 | 154,624,957 | 160,218,965 |
| Total cells | 5,777 | 8,310 | 7,029 | 9,638 |
| Median reads per cell | 22,881 | 14,739.5 | 19,083 | 14,085 |
| Total reads used in analysis | 130,047,844 | 136,742,207 | 148,241,619 | 156,671,882 |
| Cells used in analysis | 5,406 | 7,879 | 6,695 | 9,159 |

**Table S2.** Number and percentage of cells in each cluster.

| Cluster | Classification | Dataset |  |  |  |  |  |  |  |
| --- | --- | --- | --- | --- | --- | --- | --- | --- | --- |
|  |  | Uninfected and latent |  | Lytic + vehicle |  | Lytic + IDN-6556 |  | Lytic + IDN-6556 + anti-IFNs |  |
| 1 | uninfected | 2365 | 43.7% | 378 | 4.8% | 278 | 4.2% | 251 | 2.7% |
| 10 | latent | 16 | 0.3% | 387 | 4.9% | 439 | 6.6% | 420 | 4.6% |
| 13 | latent | 185 | 3.4% | 40 | 0.5% | 77 | 1.2% | 79 | 0.9% |
| 2 | latent | 1435 | 26.5% | 390 | 4.9% | 429 | 6.4% | 457 | 5.0% |
| 6 | latent (dividing) | 255 | 4.7% | 663 | 8.4% | 436 | 6.5% | 437 | 4.8% |
| 4 | latent (dividing) | 387 | 7.2% | 942 | 12.0% | 555 | 8.3% | 624 | 6.8% |
| 3 | latent (dividing) | 350 | 6.5% | 1004 | 12.7% | 499 | 7.5% | 831 | 9.1% |
| 0 | lytic (early) | 3 | 0.1% | 1829 | 23.2% | 1645 | 24.6% | 2416 | 26.4% |
| 5 | lytic (early) | 231 | 4.3% | 559 | 7.1% | 517 | 7.7% | 523 | 5.7% |
| 8 | lytic (intermediate) | 156 | 2.9% | 667 | 8.5% | 306 | 4.6% | 453 | 4.9% |
| 7 | lytic (intermediate) | 2 | 0.0% | 555 | 7.0% | 341 | 5.1% | 693 | 7.6% |
| 12 | lytic (late) | 0 | 0.0% | 412 | 5.2% | 274 | 4.1% | 456 | 5.0% |
| 9 | lytic (early) and IFN+ | 7 | 0.1% | 35 | 0.4% | 385 | 5.8% | 849 | 9.3% |
| 11 | lytic (early) and IFN+ | 14 | 0.3% | 18 | 0.2% | 514 | 7.7% | 670 | 7.3% |
|  | total | 5406 |  | 7879 |  | 6695 |  | 9159 |  |

**Table S3.**
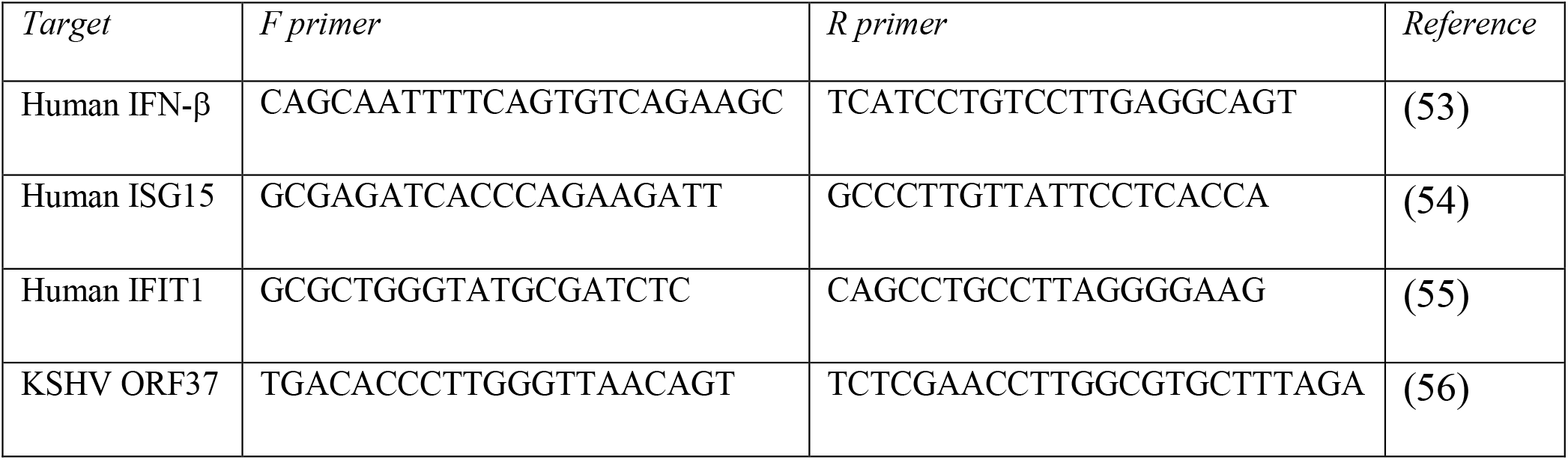
Primers used for RT-qPCR.

